# The cerebellum encodes and influences the initiation and termination of discontinuous movements

**DOI:** 10.1101/2021.06.24.449622

**Authors:** Michael A. Gaffield, Jason M. Christie

## Abstract

The cerebellum is hypothesized to represent timing information important for organizing salient motor events during periodically performed discontinuous movements. To provide functional evidence validating this idea, we measured and manipulated Purkinje cell (PC) activity in the lateral cerebellum of mice trained to volitionally elicit periodic bouts of stereotyped licking for regularly allocated water rewards. Overall, PC simple spiking modulated during task performance, ramping prior to both lick-bout initiation and termination, two important motor events delimiting movement cycles. The ramping onset occurred earlier for the initiation of un-cued exploratory licking that anticipated water availability relative to licking that was reactive to water allocation, suggesting that the cerebellum is engaged differently depending on the movement context. In a subpopulation of PCs, climbing-fiber-evoked responses also increased during lick-bout initiation, but not termination, highlighting differences in how cerebellar input pathways represent task-related information. Optogenetic perturbation of PC activity disrupted the behavior in both initiating and terminating licking bouts and reduced the ability of animals to finely time predictive action around reward delivery, confirming a causative role in movement organization. Together, these results substantiate that the cerebellum contributes to the control of explicitly timed repeated motor actions.

## Introduction

Voluntary movement often encompasses repeated starts and stops of the same deliberate motor action enabling animals to achieve their goals. Because these discontinuous movements require a temporal structure, the brain is thought to generate a timing representation of salient motor events, such as the transition to movement initiation and/or termination, that improves the consistency of periodically performed behaviors (Ivry et al., 2002). This activity may assist in the preparation for voluntary movement, as goal-directed actions must be planned prior to initiation (Ghez et al., 1991). Interconnected brain regions, including the motor cortex, thalamus, basal ganglia, and cerebellum, play a role in planning, executing, and evaluating deliberate movements (Gao et al., 2018; Guo et al., 2015; Kunimatsu et al., 2018). Human studies have helped to elucidate the putative role of each brain structure in motor control. For example, patients with cerebellar damage often have difficulty in accurately timing the initiation and termination of periodically performed discontinuous movements but can otherwise execute continuous rhythmic patterns of motor output in a relatively unimpaired manner (Bo et al., 2008; Schlerf et al., 2007; Spencer et al., 2003). These findings lend support to the idea that the cerebellum processes predictive information related to immediately impending transitions to motor action and inaction such that planned movements are finely timed and thus well executed (Bareš et al., 2019; Ivry et al., 2002; Tanaka et al., 2021). Yet, experimental validation of this hypothesis is lacking.

The cerebellar contribution to motor-action timing in the domain of sensorimotor prediction is frequently studied in animal models based on simple cue-evoked reflexive behaviors. For delay eyeblink conditioning, cerebellar activity begins ramping in response to sensory cues that predict an immediately impending conditioned response (Giovannucci et al., 2017). Similar ramping of cerebellar activity occurs during other learned behaviors in which sensory cues provide a trigger for movement initiation (Bina et al., 2020; Tsutsumi et al., 2020; Yamada et al., 2019). Neural activity also ramps in the cerebellum during motor planning tasks in which a delay period proceeds an impending deliberate movement (Chabrol et al., 2019; Gao et al., 2018; Wagner et al., 2019). As the delay time increases, the onset time of ramping activity shifts, with ramping activity commencing immediately prior to initiation, rather than occurring during the total duration of the delay (Kunimatsu et al., 2018; Ohmae et al., 2017). This ramping activity is influential in shaping behavior because its disruption can affect motor timing (Ohmae et al., 2017). Importantly, in many of these sensory-cue-driven tasks, during the delay period, animals must actively restrain themselves from initiating the motor plan too soon because false starts are often punished. Overall, the representation of both sensory cues and negative valence signals in cerebellar neural activity presents challenges in directly assessing how this brain region encodes and directly influences the timing of salient motor event transitions.

To isolate motor-event-related neural activity in the mouse cerebellum, we trained mice to perform a periodic, discontinuous movement task requiring them to elicit a stereotyped behavior at a regular interval to acquire a reward. Movement-timing activity was apparent in Purkinje cells (PCs) located in the Crus I and II lobules of the lateral cerebellar cortices. At the population level, PC simple spike firing ramped immediately before the initiation of each cycle of motor action. The ramping onset times were earlier when the movement was internally triggered compared with cases in which the same action was elicited by a sensory cue. PC simple spiking also ramped immediately before the termination of each action cycle. By contrast, climbing-fiber-driven PC activity increased only prior to movement initiation. Optogenetic perturbation of PC activity terminated ongoing movement, and movement initiated when the optogenetic stimulus ended. These perturbations impaired the ability of mice to finely time their actions around a predicted reward. Thus, the cerebellum plays an active role in the temporal organization of periodically performed discontinuous movements under volitional control.

## Results

### Mice learn to perform a discontinuous movement task based on internal timing

To understand how cerebellar activity relates to the organization of well-timed transitions to motor action and inaction during discontinuous, periodically performed movements, we trained head-fixed mice to self-initiate regular bouts of licking. For this purpose, we used an interval timing task in which thirsty mice consumed water droplets dispensed at a fixed time interval (Toda et al., 2017). In the task structure (*Figure 1A*), we randomly withheld water allocation in 20% of the trials. Because the mice voluntarily initiated licking bouts without any sensory cues during these unrewarded trials, this step allowed us to assess their ability to elicit an internally planned, periodic motor behavior that anticipated the regular timing of water-reward availability.

**Figure 1.**
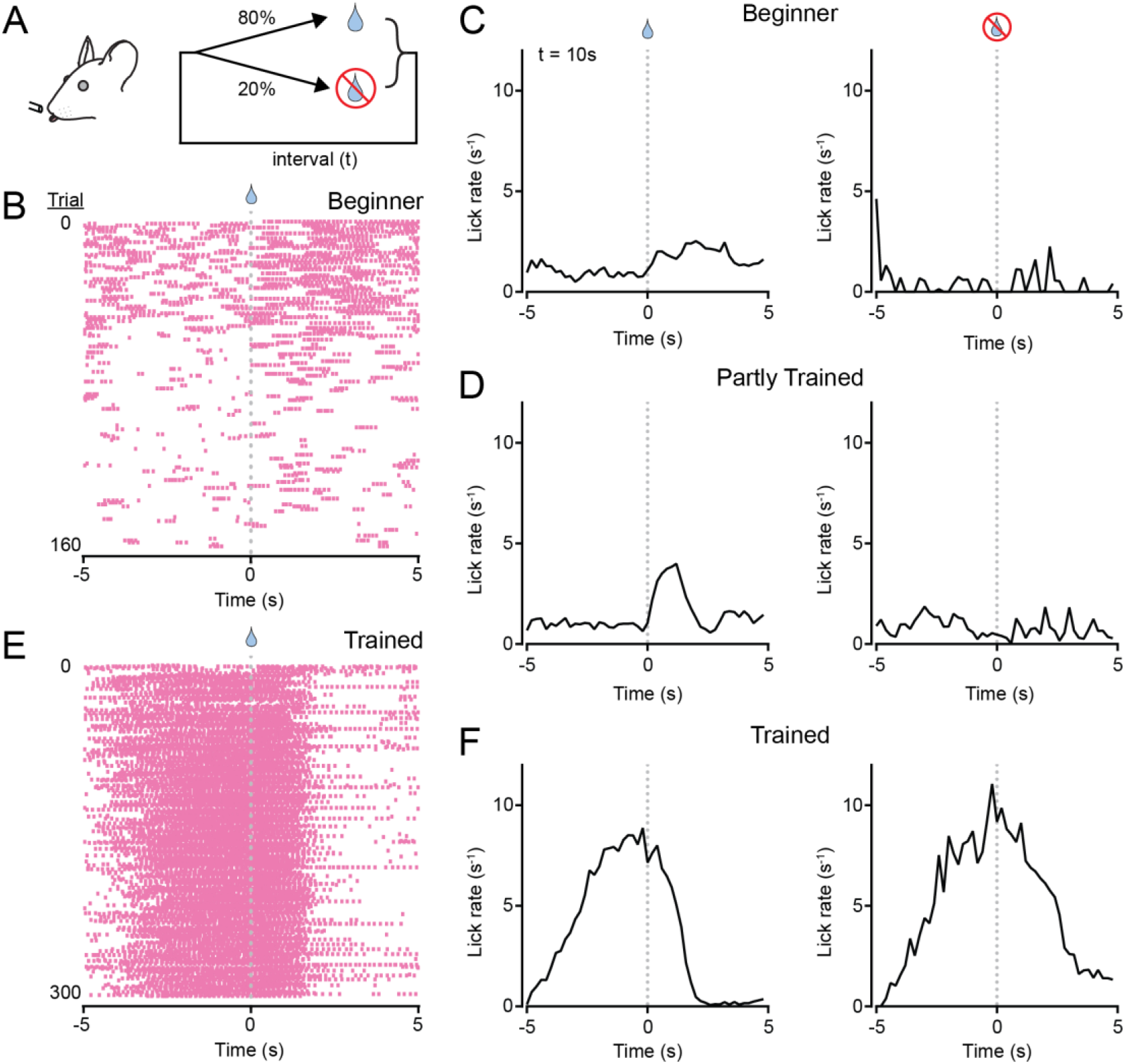
An interval timing task to assess the role of the cerebellum in organizing periodic, discontinuous movement. A. Schematic diagram of the task. Mice were trained to lick for water rewards delivered at a regular time interval (*t*). Water was allocated in most trials and withheld in the others. B. Lick patterns of a beginner mouse during an early training session. Licks are indicated by pink tic marks; water was allocated at the time indicated by the droplet (*t* = 10 s). C. Trial-averaged lick rates for the beginner mouse with session trials (159 total) separated based on whether water was allocated (left; rewarded) or withheld (right; unrewarded). D. Same as panel C but after the mouse received additional sessions of training (216 total trials). E. Lick patterns of the same animal over the course of a session after full training (same mouse as in panels B and C). F. Trial-averaged lick rates of the fully trained animal during an individual session (300 total trials).

In the first few sessions of task performance (fixed time interval of 10 s), beginner mice typically licked sporadically throughout each trial without regard to the timing of water allocation *(Figure 1B,C*). With experience, the mice learned to alter their strategy to concentrate licking bouts in response to water delivery (*Figure 1D*). Trained mice eventually initiated exploratory licking bouts prior to water-reward delivery and terminated licking shortly after consuming the dispensed water droplet (*Figure 1E,F*). This behavioral change occurred without an overt punishment to actively suppress licking during the delay period. In trained mice, the licking behavior was essentially unchanged during water omission trials, demonstrating their ability to anticipate reward timing and withhold their behavior until the next trial (*Figure 1F*). These results show that the mice reliably self-initiate regular bouts of voluntary licking based solely on internal timing, presumably referenced by the amount of elapsed time since the previous water reward, and abruptly stop licking after consuming the water reward for each trial (Rossi et al., 2016; Toda et al., 2017).

### Engagement of the cerebellum during internally timed discontinuous movements

To identify neural correlates of task-related activity in the cerebellar cortex, we used extracellular electrophysiology to record from cells in the lobules of the left Crus I and II (*Figure 2A*), regions of the lateral cerebellum implicated in orofacial behaviors (Bryant et al., 2010; Welsh et al., 1995). Although neuronal population activity was densely sampled using multi-electrode silicon probes, we restricted our analysis to PCs because they form the sole channel of output from the cerebellar cortex. We identified putative PCs based on their location, firing characteristics, and size (see Methods; *Figure 2-figure supplement 1*) (Tsutsumi et al., 2020).

**Figure 2.**
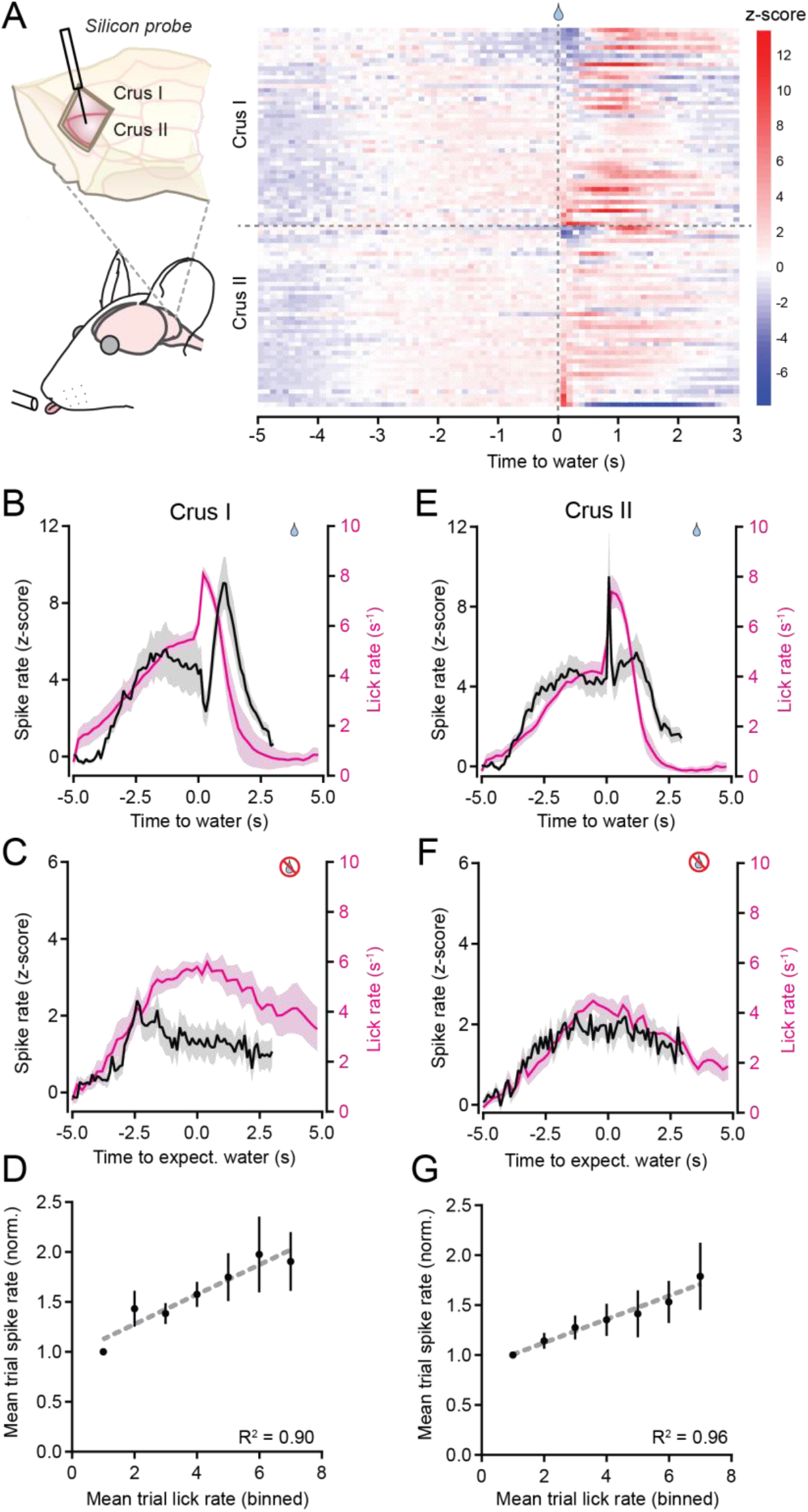
Modulation of PC simple spiking during the performance of discontinuous movement. A. Left: Electrophysiological activity was recorded from PCs using silicon probes targeting either the left Crus I or Crus II. Right: Simple spiking activity for all PCs during water-rewarded trials. Data are separated by lobule and sorted based on average activity levels within ±200 ms of water allocation (n = 47 Crus I PCs from 6 mice; n = 42 Crus II PCs from 5 mice). B. Mean simple spike rate (black) for Crus I PCs during water-rewarded trials. The corresponding trial-averaged lick rate is also shown (pink). C. Same as panel B but for trials in which water was withheld. D. Relationship between spiking activity and movement for Crus I PCs. Simple spike firing rates are binned based on the corresponding mean lick rate during the trial. Dashed line is the linear correlation. E-G. Same as panels B-D but for Crus II PCs.

In trained mice, PCs in both lobules displayed a heterogeneous range of activity changes in simple spiking patterns during task performance (*Figure 2A*). In water-rewarded trials, the average activity of Crus I PCs increased as mice began exploratory licking in anticipation of water delivery. The mean simple spike rate increased further once the mice detected water and began consummatory licking (*Figure 2B*). In unrewarded trials, the PC population activity showed a similar increase during exploratory licking (*Figure 2C*). However, in these trials, PC simple spiking lacked a prominent second peak in activity. Therefore, the uptick in PC simple spiking during consummatory licking, relative to exploratory licking, may be attributable to the encoding of reward acquisition, which has been shown to be represented in the activity of granule cells (Wagner et al., 2017). Yet, the overall simple spiking rate showed a linear correspondence to the licking rate when assessed across all contexts of the task (*Figure 2D*). Therefore, the elevation of PC simple spike firing in response to reward allocation may also reflect the abrupt increase in licking rate during water consumption (*Figure 2B*).

The average activity pattern of Crus II PCs closely resembled that of Crus I PCs. The simple spiking rate increased during exploratory licking and evolved with a further sharp uptick as consummatory licking commenced (*Figure 2E,F*). The linear relationship between the simple spiking and licking rates (*Figure 2G*) indicates that Crus II PCs also form a representation of movement. Based on these results, we conclude that PCs in both Crus I and Crus II are similarly engaged during the performance of internally timed bouts of voluntary licking. However, to more specifically determine whether PC activity in these regions encodes transitions that delimit discontinuous periodic movements, we refined our analysis to examine simple spike firing during two salient motor events: lick-bout initiation and lick-bout termination.

### PC simple spiking modulates during the transition to movement initiation

For voluntary movements, motor plans are converted into motor actions at the time point of initiation. In our task, this transition occurred at lick-bout onset, when mice began repeatedly licking at a rhythm of 6-8 Hz (Horowitz et al., 1977). To examine the simple spiking pattern at the timepoint of movement initiation, we aligned PC activity to the first lick of each bout in individual trials across animals (*Figure 3A*). Generally, the simple spiking rate began to increase several hundred milliseconds before the detection of the first lick in a bout (*Figure 3B*). Because lick initiation is rapid – the time between any visible mouth movement to full tongue protrusion is 30-60 ms (Bollu et al., 2021; Gaffield and Christie, 2017) – this ramping is more suitable for reflecting preparatory activity rather than an online representation of movement execution.

**Figure 3.**
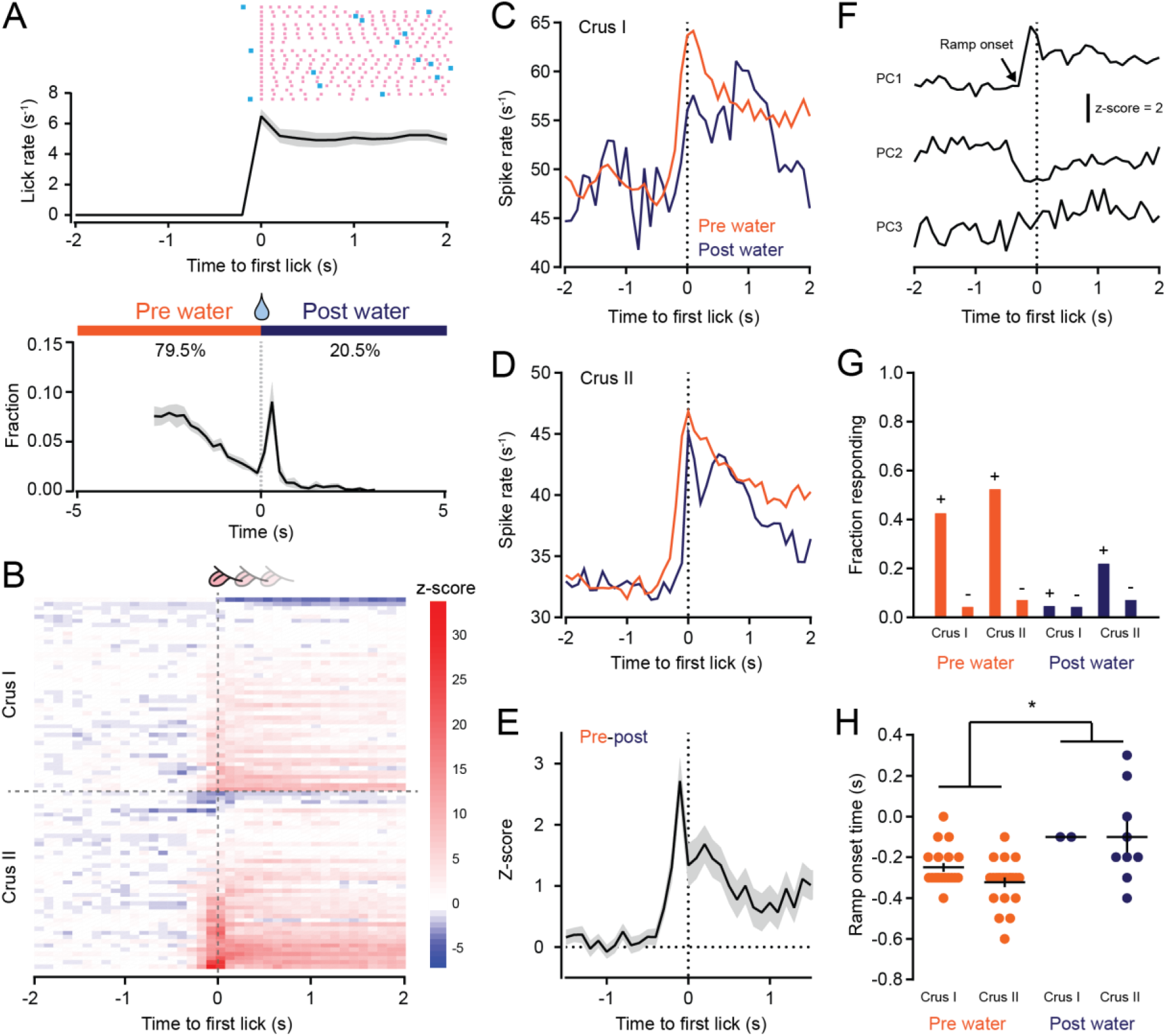
Modulation of PC simple spiking during the initiation of discontinuous movement. A. Top: plot of the mean lick rate aligned to the time point of the first lick in each water-rewarded trial bout (n = 3,164 trials from 11 mice). The lick patterns for example trials of an individual mouse are also shown with licks indicated by pink tic marks and the time of water allocation indicated in blue. Bottom: histogram of lick-bout initiation times relative to water allocation. For most trials, mice began exploratory licking prior to water delivery (pre water). However, in the remaining trials, mice refrained from licking until after water allocation (post water), which immediately triggered a rapid increase in lick-bout initiations to consume the dispensed droplet. B. Simple spiking activity for individual PCs sorted based on their average activity levels within ±200 ms of the first lick in each water-rewarded trial bout. C,D. Mean simple spike rates for Crus I (panel C) and Crus II (panel D) PCs for trials separated depending whether mice initiated lick bouts before or after water allocation (pre and post water, respectively). Note the ramps in activity prior to licking. E. Difference between the PC simple spiking rates for the two contexts of licking, calculated as a z-score. F. Individual PCs were sorted based on how their activity changed around the time of lick-bout initiation (time window of −1 to 0.5 s; see Methods); PC1 was positively modulated, PC2 was negatively modulated, and PC3 was unchanged. G. Fraction of individual Crus I and Crus II PCs with activity profiles that were either positively (+) or negatively (-) modulated around the time of lick-bout initiation, separated depending on whether licking began before or after water allocation (pre and post water, respectively). H. Comparison of the onset times of activity ramping (marked by arrow in panel F), relative to the first lick in trial bouts, for PCs whose activity positively modulated around lick-bout initiation. Data from each lobule were grouped together for statistical comparison (pre water: n = 42 PCs from 10 mice; post water: n = 11 PCs from 7 mice). Asterisk indicates significance (p = 0.0129, Student’s t-test).

Notably, the precise timing of lick-bout initiations, relative to water availability, varied from trial to trial (*Figure 3A*). As expected for trained mice that accurately anticipate impending rewards, most licking bouts were exploratory, beginning prior to water allocation (*Figure 3A*). However, in some trials, the mice did not perform any exploratory licking and, instead, waited until water became available before immediately commencing a bout of consummatory licking (*Figure 3A*). It is unclear why the mice withheld their licking until after water delivery during these trials. However, the tight distribution of lick-bout initiations around the time of water allocation in these trials (median response time: 360 ± 110 ms) indicates that the ensuing consummatory movements were reactive and were likely triggered by sensory evidence indicating water availability (i.e., the mice elicited a bout of licking during these trials only after they detected the presence of water).

Separating trials of PC activity based on whether licking bouts were initiated before or after water allocation led to an unexpected result. For PCs in both Crus I and Crus II, the average simple spiking rate began to ramp earlier for exploratory licking trials, when the movements were initiated prior to water allocation, compared with that for trials in which consummatory licking commenced immediately after water became available (*Figure 3C,D*). Across all PCs, we observed a clear temporal difference in ramping activity between these licking contexts (*Figure 3E*). These results indicate a robust activation of the PC ensemble in the lateral cerebellum prior to lick-bout initiation. However, there was clear heterogeneity in the simple spike response patterns of individual PCs. Some PCs positively modulated their firing during lick-bout initiations relative to their baseline firing rate, whereas other PCs negatively modulated their firing; the few remaining PCs were unresponsive during this motor transition (*Figure 3F*). As expected from the population average, PCs that positively modulated their simple spike activity were much more common than PCs that negatively modulated their activity (*Figure 3G*). Additionally, PCs were more likely to positively modulate their activity (i.e., ramp) prior to exploratory licking bouts that preceded water allocation than for bouts of consummatory licking that were reactive to water allocation (*Figure 3G*). In the subset of PCs with positively modulated activity, a comparison of ramping onset times across conditions revealed that activity began ~200 ms earlier for bouts of exploratory licking compared with bouts of purely reactive consummatory licking (*Figure 3H*). We conclude that PC simple spike rates change prior to the initiation of licking bouts, with the timing of ramping activity depending on whether the ensuing movement was initiated by internal motivation or was triggered by a sensory cue indicating water availability.

### PC simple spiking modulates during the transition to movement termination

The completion of a motor action is also a salient event. Therefore, to investigate whether PCs encode information pertaining to movement termination, we aligned PC simple spike activity to the last lick of a lick bout (*Figure 4A,B*). PCs displayed heterogeneous output patterns during this motor transition, with some cells positively modulating their activity and others negatively modulating their activity (Figure 4B). On average, the simple spiking rate in the PC population abruptly increased just prior to the last lick for cells in both Crus I and Crus II during water-rewarded trials (*Figure 4C*). Because the last lick in a bout occurred well after the time point of water delivery (median time interval: 1.37 ± 0.21 s; *Figure 4D*), most of this ramping activity occurred after the dispensed water droplet had been largely consumed. In addition to encoding the impending termination of a lick bout, this simple spiking increase may also represent information pertaining to swallowing and/or reward signaling by satiation in the gastrointestinal system (Augustine et al., 2019; Zimmerman et al., 2019). However, simple spiking also ramped prior to the last lick in bouts elicited during water omission trials (*Figure 4E*). Thus, the ramping activity in PCs at the end of licking bouts most likely corresponds to the anticipation of motor-action termination. Overall, lick-bout termination was well represented in the PC activity in both lobules (36.2% and 23.8% of PCs in Crus I and Crus II, respectively, modulated their simple spiking [see Methods]).

**Figure 4.**
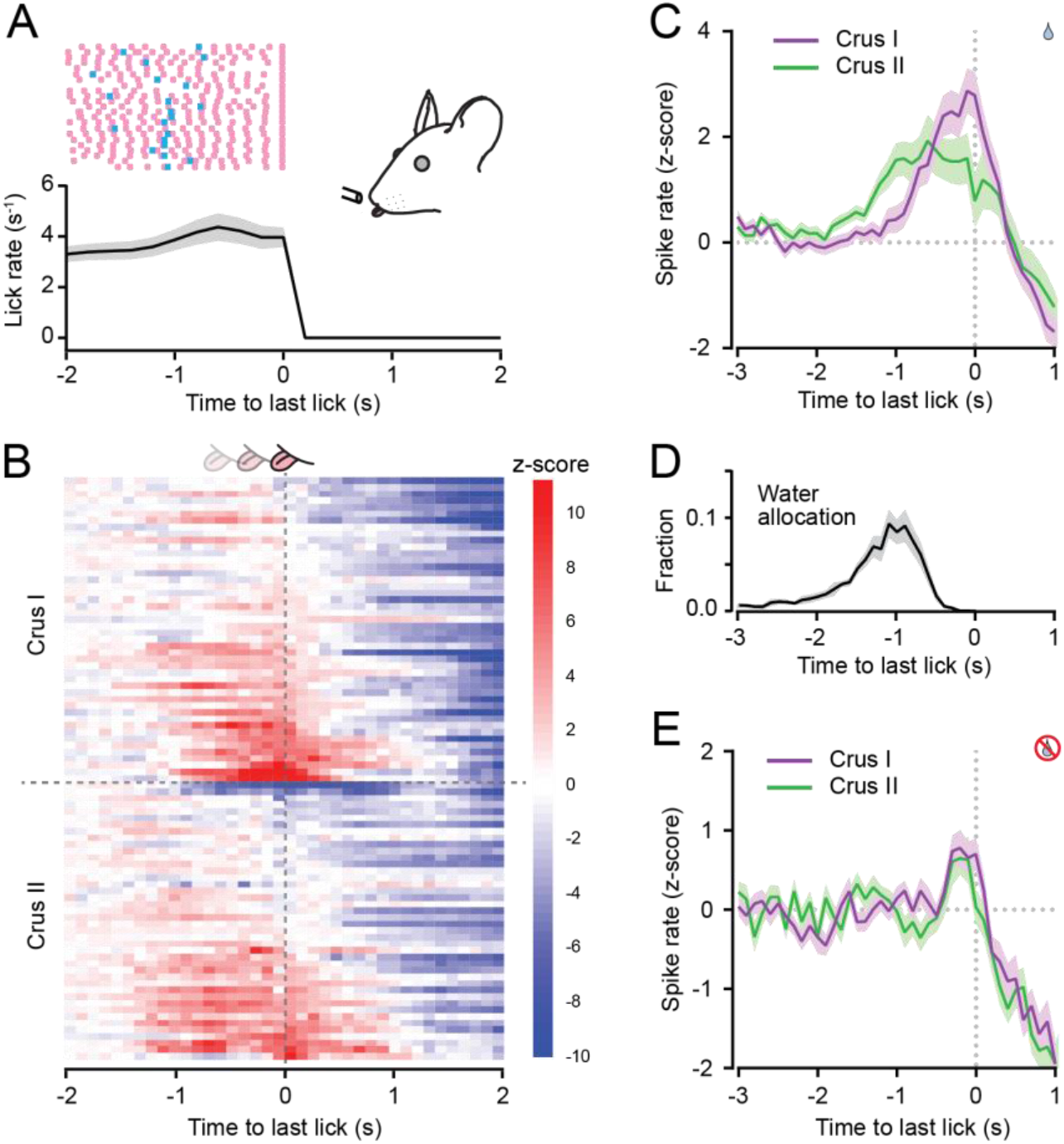
Modulation of PC simple spiking during the termination of discontinuous movement. A. Plot of mean lick rate aligned to the last lick in water-rewarded trial bouts (n = 2,846 trials from 11 mice). Also shown are lick patterns of example trials for an individual mouse with licks indicated by pink tick marks and the timing of water allocation in blue. B. Simple spike activity for individual PCs sorted based on their average activity levels within ±200 ms of the last lick in each water-rewarded trial bout. C. Trial-averaged simple spike activity for Crus I (n = 47 from 6 mice) and Crus II (n = 42 from 5 mice) PCs aligned to the time of the last lick in water-rewarded trials. D. Distribution of the timing of water allocation aligned to the point of the last lick in rewarded trial bouts (same trials as in panel A). E. Same as panel C but for unrewarded trials (n = 44 Crus I PCs from 6 mice; n = 38 Crus II PCs from 5 mice).

To examine whether individual PCs tune their activity to specific task features, we sorted all cells based on how their simple spiking modulated during motor-event transitions. PCs were categorized whether they exhibited ramp firing to exploratory licking initiated prior to water allocation, ramp firing to consummatory licking initiated after water allocation, and/or ramp firing at the end of either type of licking bout. Because negatively modulated responses were relatively rare, we pooled PCs that exhibited decreased firing during any motor transition into a single group. Plots of mean spiking rates from several of these groupings, including PCs that positively modulated their firing around either the first or last lick in a bout (*Figure 5A,5B*) or that negatively modulated their firing (*Figure 5C*), confirmed that our sorting differentiated PCs based on their response profiles. Overall, individual PCs in both Crus I and Crus II were heterogeneous in their representation of task-related attributes (*Figure 5D*). Although specialist PCs were common, for example, showing a preference for ramp firing at the initiation of exploratory licking, many PCs were engaged by multiple types of motor transitions, e.g., by increasing their firing to both licking initiation and termination. In summary, the simple spiking activity of PCs in the lateral cerebellar cortex modulates in response to salient motor events during discontinuous bouts of periodic movements, with only modest tuning for a specific type of motor transition.

**Figure 5.**
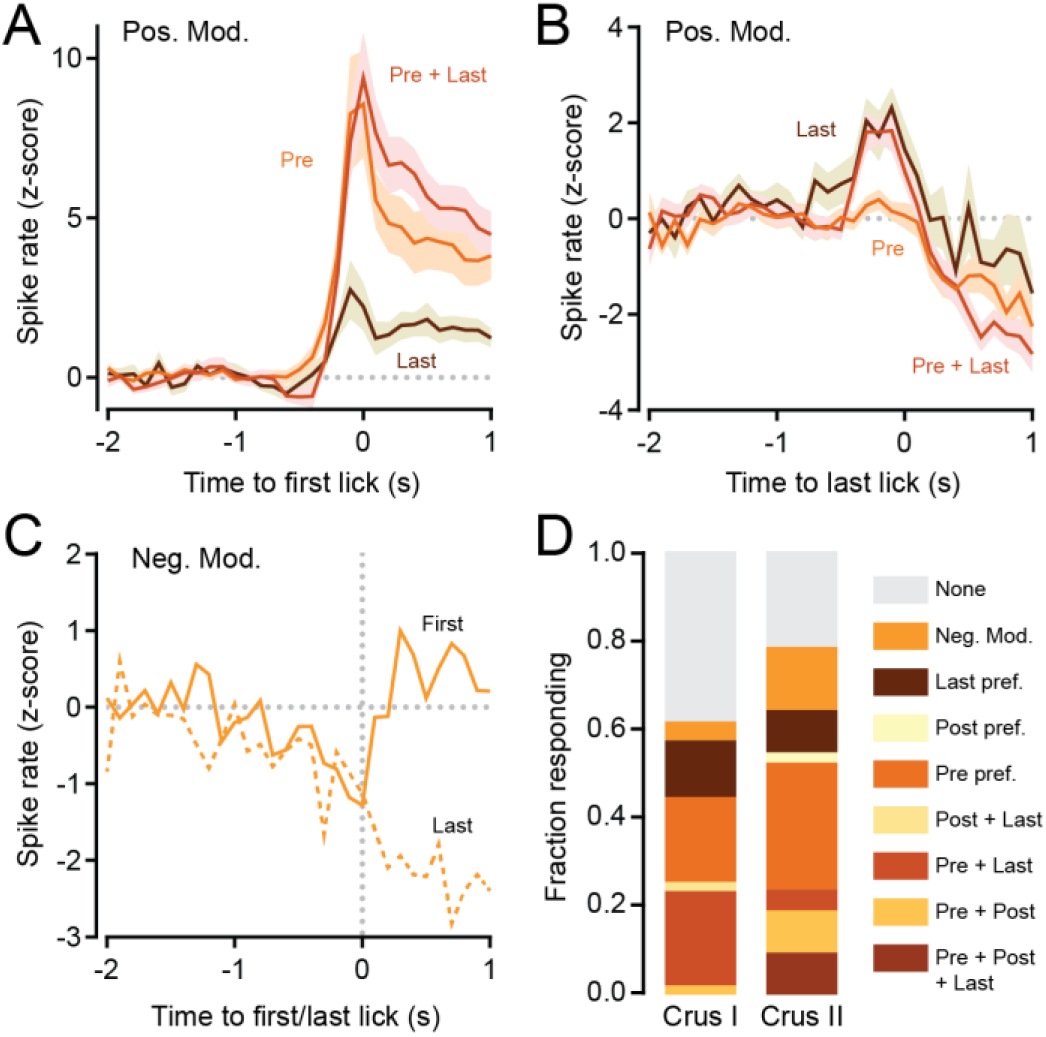
Task-feature tuning of individual PCs. A. Trial-averaged activity profiles of PCs grouped based on the preference of their simple spiking profile for specific motor events during the task. PCs characterized as having a biased increase in firing to the first lick of exploratory bouts (Pre, n = 21 PCs from 9 mice) or to both the first licks of exploratory bouts as well as the last licks (Pre + Last, n = 16 PCs from 5 mice) showed a robust change at lick bout initiation. By contrast, PCs characterized as preferentially responding only to the last lick in trial bouts (Last, n = 5 PCs from 3 mice) had a much smaller increase in simple spiking at lick bout initiation. B. Same PCs as panel A but during the time around the last lick in trial bouts. For last-lick-preferring PCs, as well as those that responded to both the first and last licks, there was a robust increase in firing at lick bout termination. However, PCs with biased firing for only the first lick in exploratory bouts did not show any activity changes around lick bout termination. C. Trial-averaged simple spiking profiles of PCs characterized as having negatively modulating responses during first licks (solid line, n = 7 PCs from 6 mice) or last licks (dashed line, n = 5 PCs from 3 mice) during trial bouts. SEM ranges are removed for clarity. D. Summary of tuning profiles for Crus I and Crus II PCs (n = 47 and 42, respectively). PCs with a decrease in firing (Neg. Mod.) during either the first or last licks in trial bouts were considered as a single group as well as those without any discernable activity tuning (None).

### Climbing-fiber-induced PC activity increases during movement initiation

In addition to simple spikes, PCs also fire complex spikes and simultaneous bursts of dendrite-wide calcium action potentials in response to excitation provided by climbing fibers, the axonal projections of inferior olive neurons (Llinas and Sugimori, 1980; Ozden et al., 2009). Therefore, to determine the representation of climbing-fiber-induced PC activity during periodically performed discontinuous movements, we used two-photon imaging to measure climbing-fiber-evoked dendritic calcium events in PCs expressing the calcium sensor GCaMP6f (*Figure 6A*). This imaging-based approach is more sensitive in detecting climbing-fiber-evoked activity because it is challenging to reliably distinguish complex spike waveforms in extracellular electrophysiological unit recordings (Sedaghat-Nejad et al., 2021; Tsutsumi et al., 2020). In quiescent mice, individual calcium events were readily apparent in the dendrites of left Crus I and II PCs (*Figure 6B*), reflecting that climbing fibers continuously bombard PCs at 1-2 Hz (Gaffield et al., 2016; Mukamel et al., 2009; Ozden et al., 2012). However, during task performance, only a subset of the PCs showed a behavior-induced change in activity. This evoked response appeared to be spatially organized and dependent on the behavioral context. In water-rewarded trials, the average rate of climbing-fiber-evoked dendritic calcium events increased in PCs in some imaged regions of Crus I and Crus II around the time of water allocation. However, in other imaged regions, there was no change in average PC calcium activity (*Figure 6C*). Interestingly, the increase in climbing-fiber-evoked activity during water consumption was absent in the same PCs during water omission trials, when mice elicited licking bouts but did not receive water rewards (*Figure 6D*). Thus, climbing fibers appear to signal reward-acquisition-related information to PCs in specific regions of Crus I and Crus II (Heffley and Hull, 2019; Heffley et al., 2018; Kostadinov et al., 2019).

**Figure 6.**
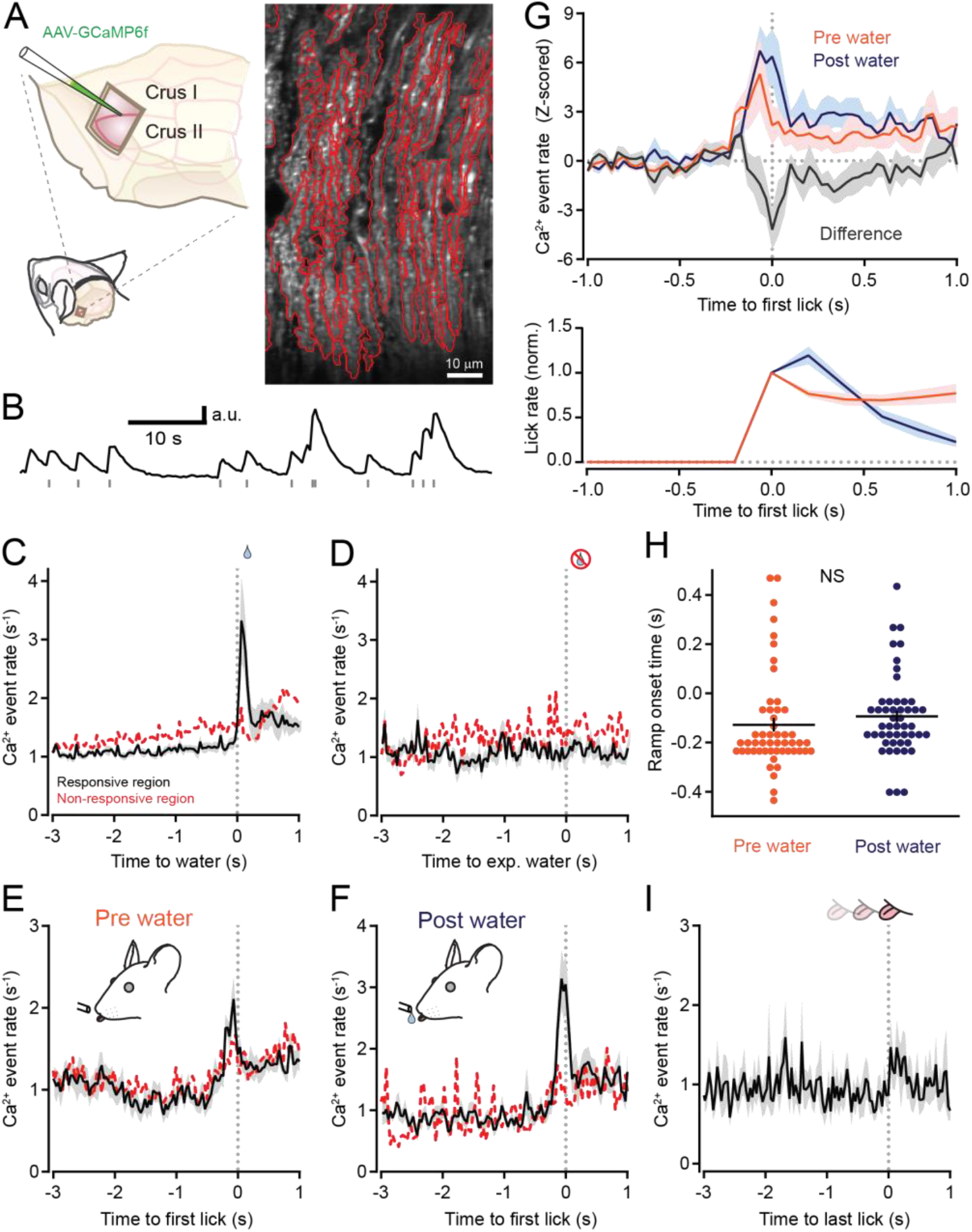
Climbing-fiber-evoked PC activity increases at the initiation of discontinuous movement. A. Left: AAV containing GCaMP6f under control of the *Pcp2* promoter was injected into Crus I and Crus II to specifically transduce PCs; a cranial window provided optical access to the infected region. Right: A two-photon image with identified PC dendrites outlined in red. B. Example fluorescence trace from a PC dendrite showing spontaneous calcium activity during quiescence. Individual climbing-fiber-evoked calcium events are indicated by gray tic marks. C. Average climbing-fiber-evoked calcium event rates aligned to the time point of water delivery for water-rewarded trials. PC dendrites in some regions of Crus I and Crus II showed clear increases in activity when mice elicited bouts of licking to water allocation (black line, n = 377 PCs in 8 ROIs from 4 mice), whereas in other regions there was very little to no change in activity (dashed red line, n = 239 PCs in 5 ROIs from 4 mice). D. Same as panel C but for licking during unrewarded, water-omission trials. E. Trial-averaged calcium event rates in PC dendrites aligned to the timing of the first lick in exploratory bouts initiated prior to water allocation (same data as panel C). F. Same as panel E but aligned to the first lick for bouts of consummatory licking initiated after water allocation. G. Top: Overlay of trial-averaged calcium event rates for PCs in task-responsive regions of Crus I and II, aligned to the first lick of bouts initiated before or after water allocation (pre and post water, respectively). The subtracted difference is also shown. Bottom: Plot of the corresponding normalized lick rate for these trial types (n = 8 sessions from 5 mice). Note the uptick in licking rate after the first lick for consummatory bouts initiated following water allocation. H. Comparison of onset times for climbing-fiber-evoked calcium event ramping for individual PCs in trials where licking was initiated before (pre water; n = 52 PCs) or after (post water; n = 49 PCs) water allocation (see Methods). Black line shows the mean (not significant, NS; p = 0.36, Student’s t-test). I. Trial-averaged calcium event rate aligned to the timing of the last lick in trial bouts (n = 616 PCs, n = 12 sessions, 5 mice).

To more carefully evaluate the correspondence of climbing-fiber-evoked activity in PCs around motor event transitions, we aligned dendritic calcium activity to the first licks in bouts of either exploratory licking initiated prior to water allocation or consummatory licking initiated after water delivery. Average calcium event rates ramped prior to movement initiation for both licking contexts. These increases in climbing-fiber-evoked activity were prominent in the reward-responsive regions of Crus I and Crus II (*Figure 6E,F*). Dendritic calcium event rates increased to a slightly greater extent for bouts of consummatory licking compared with exploratory licking (*Figure 6G*). This difference may result from an increased level of motor drive, as licking rates were higher during bouts of consummatory licking compared with bouts of exploratory licking (*Figure 6G*). The onset times of calcium event ramping, relative to the detection of the first lick, were similar for the initiation of both exploratory and consummatory bouts of licking (*Figure 6H*). Aligning PC dendritic calcium events to the last lick of lick bouts did not reveal any clear change in climbing-fiber-evoked activity around the transition to action completion (*Figure 6I*). Together, these results indicate that climbing-fiber-induced activity in a specific population of PCs ramps prior to the initiation, but not the termination, of both internally timed and sensory-cued bouts of goal-directed motor behavior.

### Optogenetic PC stimulation can both initiate and terminate movement

Having established that PCs in the lateral cerebellar cortex modulate their activity during the performance of discontinuous periodic movements, we applied an optogenetic approach to examine the causal role of PC activity in coordinating motor event transitions between action and inaction. We obtained conditional expression of channelrhodopsin-2 (ChR2) in all PCs by crossing transgenic Ai27 mice with the Pcp2::Cre driver line (Madisen et al., 2012; Zhang et al., 2004). In these animals, PC photostimulation drove robust simple spike firing, as measured *in vivo* using extracellular unit recording under anesthesia (*Figure 7A,B*). To optogenetically perturb PC activity during task performance, we bilaterally implanted optical fibers above the left and right Crus II lobule and introduced light stimuli during the period of peak anticipatory licking in a randomized subset of water omission trials when trained mice were robustly engaged in performing internally timed, exploratory movements without any sensory evidence indicating water availability (Figure 7C,D). In response to PC photostimulation, the licking rate slowed and became erratic, showing a sharp degradation in rhythmicity; eventually, most mice ceased licking altogether (*Figure 7D,E*). After the PC photostimulation period ended, the mice resumed consummatory licking, albeit at a diminished rate compared with control trials (*Figure 7D*). Unilateral photostimulation of PCs in either the left Crus I or Crus II lobule led to less dramatic effects on licking rate and rhythmicity (*Figure 7E* and *Figure 7-figure supplement 1A*). The light stimulus had no effect on licking in mice that did not express ChR2 (*Figure 7E*). Therefore, optogenetically disrupting PC activity during internally timed licking severely disrupts behavioral performance.

**Figure 7.**
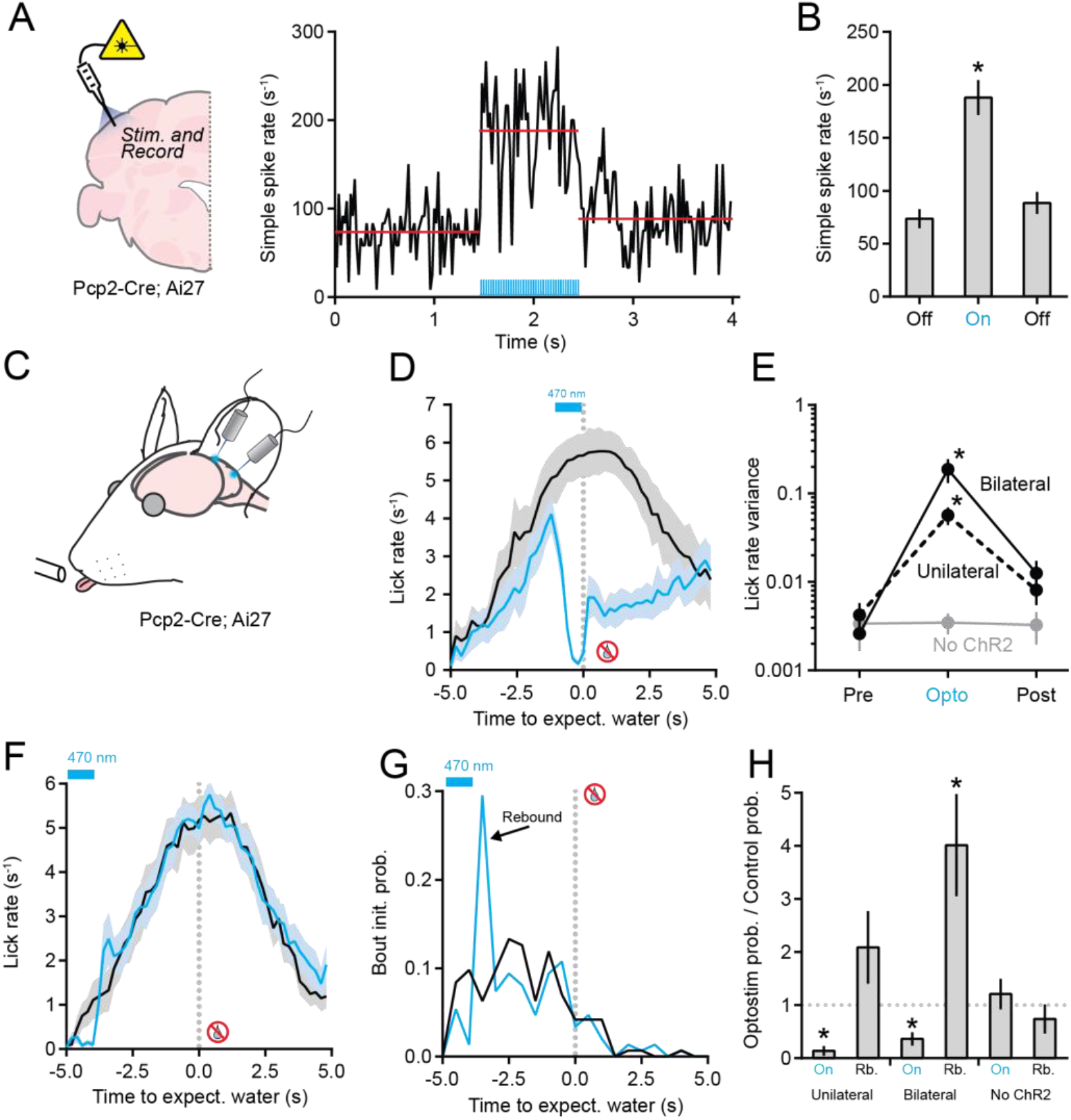
Optogenetic perturbation of PC activity degrades the performance of discontinuous movements. A. Left: Extracellular electrophysiological measurements were obtained from ChR2-expressing PCs in response to photostimulation. Right: Optogenetically induced simple spiking in a PC (light pulses indicated in blue). Red lines show the means for each epoch. B. Summary plot of mean simple spike rate across PCs (n = 6) before, during, and after the optogenetic stimulus. Asterisk indicates significance (p < 0.0001, ANOVA with Tukey’s post-test). C. In mice with ChR2-expressing PCs, Crus II was bilaterally photostimulated during the interval task. D. The effect of bilateral optogenetic PC activity perturbation on lick rate (blue) in unrewarded trials (n = 9 sessions, 3 mice). The photostimulus was timed to the period of peak licking, as referenced by interleaved control trials (black). E. Summary of lick rate variability during optogenetic perturbation of PC activity. The y-axis is scaled logarithmically. Data include trials with bilateral photostimulation of Crus II (n = 9 sessions, 3 mice), trials with unilateral photostimulation of Crus I or II (n = 12 sessions, 4 mice), and trials in control mice where blue light was delivered bilaterally to the cerebellum but PCs did not express ChR2 (n = 6 sessions, 2 mice). Asterisks indicate significant differences during photostimulation trials (p = 0.0228 and 0.0199 for the unilateral and bilateral photostimulation conditions, respectively; ANOVA with Tukey’s post-test). F. Same as panel D but with the photostimulus timed to the period of earliest lick bout initiations. Note the absence of licking during photostimulation and the large increase in licking immediately after photostimulation ended (n = 11 sessions, 3 mice). G. Histogram of lick bout initiation times for control and optogenetic stimulation trials (same data as panel F). A clear increase in licking probability is apparent after photostimulus ended. H. Summary of the effect of optogenetic PC activity perturbation on licking behavior during task performance. Asterisks indicate a significant reduction in licking during photostimulation (On: p = 0.0079 and p = 0.0486 for unilateral and bilateral stimuli, respectively; ANOVA with Tukey’s correction for multiple comparisons) and a significant increase during the rebound period for bilateral stimulation (Rb: p = 0.0034, ANOVA with Tukey’s correction for multiple comparisons). In the control condition, light stimuli were delivered to the cerebellum of non-ChR2-expressing animals (n = 12 sessions, 2 mice).

To further explore the ramifications of PC activity perturbation on motor event transitions, we delivered optogenetic stimuli well before the time point of expected water allocation, when mice first began to elicit bouts of exploratory licking. During the PC photostimulation period, exploratory licking again largely abated (*Figure 7F* and *Figure 7-figure supplement 1B*). However, immediately after the photostimulation, the mice initiated a barrage of licking at a probability much greater than that observed at the same time point in control trials (*Figure 7G,S2C*). Such optogenetically induced rebound licking was more prominent for bilateral photostimulation of Crus II PCs than for unilateral photostimulation of Crus I or Crus II PCs and was not observed for light stimuli delivered to non-ChR2-expressing mice (*Figure 7H*). Together, these results indicate that PC photostimulation can both initiate and terminate bouts of licking, depending on the timing of the activity perturbation relative to the licking context, indicating a role for PC activity in coordinating motor event transitions.

### Optogenetic perturbation of PC activity disrupts motor timing

We were surprised to find that despite the uptick in early lick initiations due to the optogenetic stimulus, the mice accurately anticipated the reward timing, showing a peak licking rate comparable to that in control trials around the time point of expected water allocation (*Figure 7F*). We hypothesized that the duration between the optogenetic PC activity perturbation and expected reward availability allowed the mice to recover the proper timing of their behavior. Therefore, we re-trained a cohort of mice and progressively shortened the interval between water allocation trials so that the mice would initiate licking closer to the time point of water availability (*Figure 8A-C*). In response to this change, the mice adjusted the timing of their licking bouts, taking only a few sessions to learn to accurately anticipate the new water delivery interval (5 s). In recordings from trained mice, Crus II PCs fired simple spikes during short-interval trials at a rate similar to that observed in long-interval trials (*Figure 8-figure supplement 1A,1B*). Likewise, bilateral optogenetic perturbation of Crus II PC activity around the time point of early lick initiations similarly disrupted behavioral performance. Mice ceased licking during the photostimulation period and elicited rebound licking bouts when the optogenetic stimulus ended (*Figure 8B*). Interestingly, for optogenetic PC perturbations during short-interval trials, there was a clear degradation in the ability of the mice to accurately anticipate reward timing: their peak licking rate was delayed relative to the control trials, occurring well after the expected time point of water allocation (*Figure 8B*).

**Figure 8.**
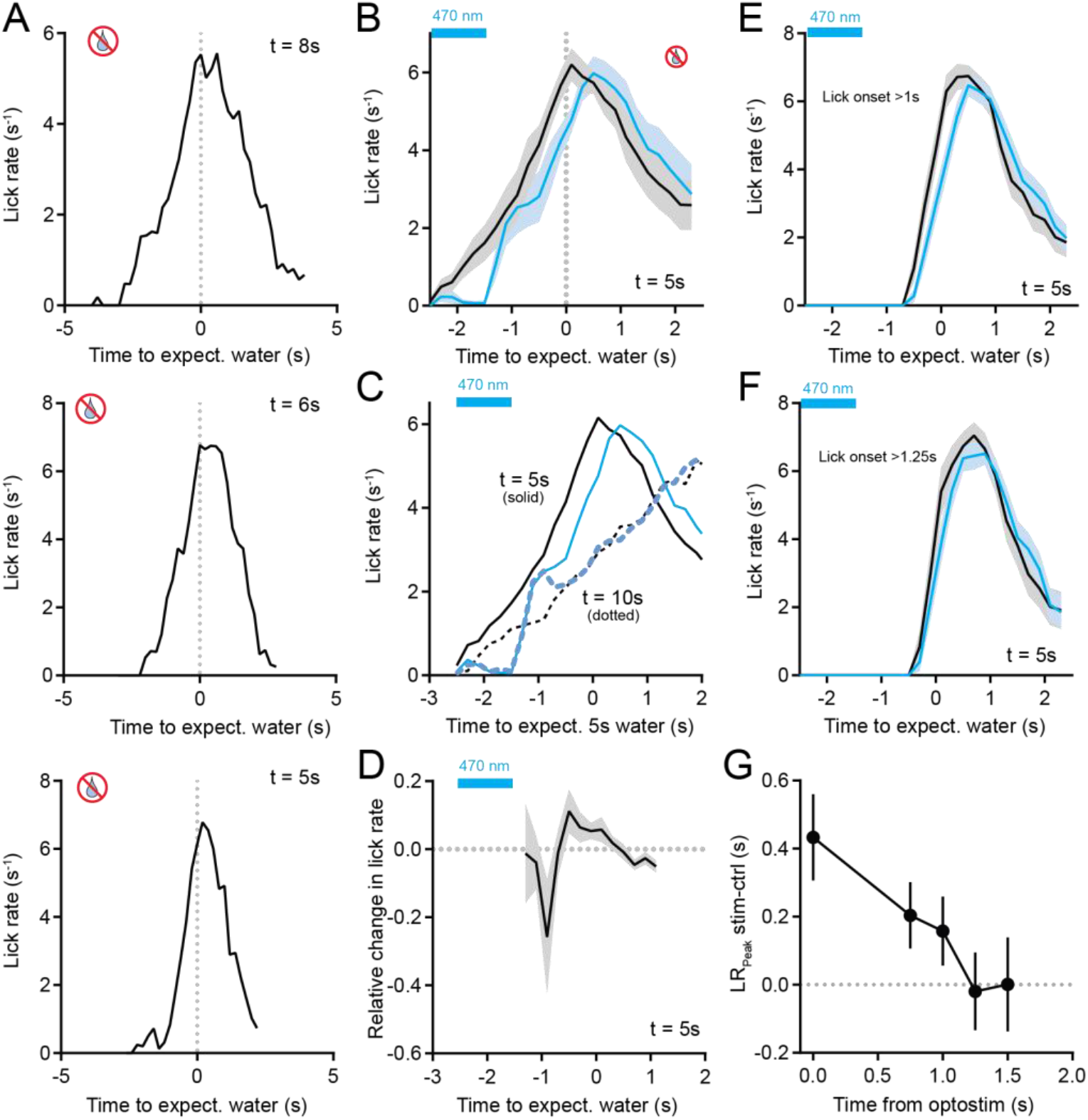
PC activity perturbation during the initiation of discontinuous movement disrupts the timing of subsequent goal-directed action. A. Example of licking activity in water-omission trials as an individual mouse progressively learned shorter time intervals of water delivery over the course of subsequent training sessions (*t* = 8, 6, and 5 s for the single sessions shown in the top, middle, and bottom panels, respectively). Note how well the mouse sharpens its response for the shortest interval around the time of expected water-reward delivery. B. The effect of bilateral optogenetic activity perturbation of Crus II PCs on licking (blue), relative to interleaved control trials (black). A shift in the timing of the peak lick rate is apparent in the photostimulation trials (n = 11 sessions, 3 mice). C. Comparison of the effect of PC activity perturbation on licking for two different water delivery intervals (*t* = 10 and 5 s, dotted and solid lines, respectively). There were nearly identical effects during, and just after, the photostimulation period. However, in contrast to the longer interval trials, licking during the shorter interval trials did not recover to control levels after the photostimulation period. D. Plot showing the relative change in lick rate induced by the optogenetic perturbation for short-interval trials (subtracted difference between the stimulation and control trials). A large reduction in lick rate was apparent for approximately 1 s after the photostimulation ended. E,F. Comparison of lick rate in control and stimulation trials, separated based on whether the animal chose to lick at least 1 s (panel E) or 1.25 s (panel F) after photostimulation. A shift in peak lick rate was evident when licking began shortly after the photostimulation ended. G. The difference in timing of the peak lick rate in stimulation trials, relative to control, binned based on the time between the optogenetic stimulus and the onset of licking.

Comparing licking behaviors across mice in response to PC photostimulation for trials of short or long reward-allocation intervals (5 and 10 s, respectively; different sessions) provided insight into when PC activity plays a dominant role in shaping the timing of anticipatory movements. During the photostimulation period, optogenetic perturbation disrupted the licking behaviors to a similar extent (*Figure 8C*). However, for short-interval trials, licking rates after the post-stimulus rebound in motor output remained below the level of session-matched control trials. In contrast, for long-interval trials, licking rates rapidly recovered to control levels (*Figure 8C*). The reduction in licking rates induced by PC activity perturbation in short-interval trials, relative to control levels, seemed to occur within a 1 s window immediately after the optogenetic stimulus period (*Figure 8D*). Therefore, we sorted short-interval trials into groups based on when the mouse initiated a licking bout relative to the end of the optogenetic stimulus. For lick bouts beginning as late as 1 s after the optogenetic stimulus, the time of peak licking was delayed compared with that in control trials (*Figure 8G*). However, this difference was nearly absent when licking bouts were initiated more than 1.25 s from the end of the optogenetic stimulus (*Figure 8H*). Across all short-interval trials, there was a clear decaying time dependence of the effectiveness of PC activity perturbation in delaying the timing of the peak anticipatory licking rate (*Figure 8I*). Therefore, within a narrow temporal window, PC activity is required at the transition to motor action for mice to accurately structure their licking to properly anticipate reward timing, emphasizing the role of the cerebellum in the fine control of discontinuous periodic movements (Ivry et al., 2002).

## Discussion

The temporal consistency of periodically performed discontinuous movement is believed to be aided by timing representations of salient motor events by the cerebellum (Ivry et al., 2002). In support of this idea, we observed changes in murine PC activity immediately prior to both the initiation and termination of licking bouts timed to regular intervals of water-reward allocation. Moreover, perturbation of this activity influenced the behavioral performance. Although the cerebellum has been implicated in learning temporal associations of predictive sensory cues and impending motor actions, the changes we observed in PC activity occurred while the animals elicited goal-directed licking independent of any external stimulus. Thus, our results indicate that this activity is internally driven and related to the timing of motor events that are pertinent to organizing the temporal consistency of volitional behavioral performance. Our results reinforce the idea that the cerebellum influences well-timed, regularly performed actions that are generated solely by motivation in addition to actions that are sensory driven.

We found that intermingled PCs in the Crus I and II lobules displayed heterogeneous coding of movement-related task parameters in their pattern of simple spiking. In our recordings, we readily observed activity ramping in individual PCs in advance of both lick-bout initiation and termination, suggesting a strong representation of impending motor-event transitions in this population. Activity ramping in PCs has also been observed during other volitional motor behaviors. For example, in monkeys, PCs fire elevated barrages of simple spikes immediately prior to changes in eye-movement speed and/or direction during object tracking (Herzfeld et al., 2015) or arm movements during reaching behaviors (reviewed in Ebner et al., 2011). Increased simple spiking also briefly precedes spontaneous and sensory evoked whisking in some PCs in mice (Brown and Raman, 2018; Chen et al., 2016). However, compared to the close temporal correspondence between PC activity ramping and motor action in these earlier reports, lasting just a few tens of milliseconds, we found that the onset time of simple spike ramping preceded lick bout initiation and termination by hundreds of milliseconds. This time frame is consistent with the ramping of cerebellar activity in advance of internally timed, volitional eye movements of monkeys that followed a delay interval of several seconds (Ohmae et al., 2017). Thus, the onset of PC activity ramping, relative to the ensuing movement, may undertake different dynamics dependent on the timing needs of the underlying behavior.

PC activity did not map the entire epoch of the delay period in our interval timing task, but rather only the end point immediately prior to lick-bout initiation. This result is consistent with the idea that multiple brain regions form timing representations of movements at different scales, with the cerebellum only contributing to the sub-second range (Tanaka et al., 2021). Interesting, we found that the onset time of PC activity ramping occurred earlier for licking that was preemptive, rather than reactive, to water-reward availability. Because reactive licking was likely triggered by an external cue, it may be that the shorter onset time of ramping in this context of licking promotes a more reflex-like response, perhaps emergent from sensorimotor associations formed in the cerebellum, ensuring that ensuing consummatory movements are executed with little delay. By contrast, when the timing representation of the impending motor event is internally rather than externally signaled, such as for exploratory licking elicited prior to water allocation, PC activity may ramp earlier due to input (either direct or indirect) from a separate brain region that also participates in organizing the behavior. Contributing brain regions could include the basal ganglia, which also forms a time representation of discontinuous movement, but with a longer timescale than the cerebellum (Kunimatsu et al., 2018; Ohmae et al., 2017). In mice, optogenetic stimulation of an inhibitory basal ganglia pathway catastrophically disrupts licking during the same interval timing task, indicating that this region can also play a role in organizing the behavior (Toda et al., 2017). The cerebellum is also recurrently connected with the cerebral cortex, forming a loop whose activity helps maintain motor plans in working memory until initiation (Gao et al., 2018; Svoboda and Li, 2018). Because mice must carefully track the passage of time to correctly anticipate their next planned cycle of licking around water-reward allocation, earlier PC activity ramping during exploratory licking bouts may also reflect the engagement of cortical circuitry.

The termination of motor action is also a salient event that delimits discontinuous movement. However, in comparison to movement initiation, less is known about how cerebellar activity at the end of each movement cycle encodes and influences periodically performed behaviors. We observed widespread ramping of simple spiking in individual Crus I and II PCs at lick bout termination. During task training, the mice learned to adjust their licking rate, ultimately stopping after consuming the dispensed water droplet on rewarded trials. Mice also stopped licking on most unrewarded trials, presumably because the estimated time window for the expected water reward had passed. Thereafter, mice began waiting for the next period of reward allocation to elicit their planned behavior. Thus, we speculate that the uptick in PC simple spiking toward the end of licking bouts prepares an impending stop to motor action, similar to the modulation of cerebellar activity at the end of dexterous reaching behaviors that influences kinematics and endpoint precision of grasps (Becker and Person, 2019; Low et al., 2018). We propose that this activity may be necessary to coordinate precise temporal control of the subsequent cycle of movement.

In addition to the ramping of PC simple spiking, we observed an increase in climbing-fiber-evoked activity in a PC subpopulation, which preceded lick-bout initiation by ~100 ms. Although climbing fibers are responsive to sensory stimuli (Gaffield et al., 2019; Ohmae and Medina, 2015), the activity increase occurred in advance of exploratory licking bouts that were initiated without an overt sensory cue, ruling out the possibility that this activity represented an external stimulus. Climbing-fiber-evoked activity did not change at lick-bout termination, suggesting a specific role for this input only at the start of each movement cycle. Reward-related climbing fiber signaling is prevalent in the cerebellar cortex (Heffley and Hull, 2019; Heffley et al., 2018; Kostadinov et al., 2019); thus, the dramatic increase in climbing-fiber-evoked activity in PCs during behavior could reflect a prospective response to an anticipated reward. However, the same PCs were unresponsive in water-omission trials, indicating a lack of apparent reward prediction errors. Therefore, we conclude that movement-aligned activity is motor-related.

Because the synchronization of climbing-fiber-evoked complex spiking within parasagittal-aligned clusters of PCs can evoke and/or invigorate motor action, including bouts of sensory-triggered licking (Apps and Hawkes, 2009; Ten Brinke et al., 2017; Tsutsumi et al., 2020; Welsh, 2002), the activity increase we observed in PCs due to climbing fiber input could influence the kinematics of lick-bout initiation if this activity were temporally correlated in the PC population. However, as we used small fields of view during our optical recordings, there were generally too few simultaneously active PC dendrites to accurately quantify the level of their synchrony. For this reason, we were unable to determine whether the responsive and unresponsive regions of Crus I and II during task performance corresponded to distinct, functional clusters of PCs that have been shown to be either engaged or not engaged during sensory-driven licking (Tsutsumi et al., 2020).

Our optogenetic experiment provided causal evidence that PC activity can elicit motor event transitions resembling those occurring during periodically performed licking. Perturbating PC activity terminated ongoing licking, even when the stimulus was timed to the peak output rate around water rewards. This perturbation could also trigger lick-bout initiations when the photostimulation ended. However, optogenetically evoked licking was apparent only when the perturbation was timed to the period when the animals were beginning to elicit exploratory bouts of licking in anticipation of water rewards. Therefore, like the susceptibility of the cerebellum to instantiate learning (Albergaria et al., 2018), our results indicate that there may be a conditional state during which the cerebellum is more effective at triggering movement initiation, in particular when planned movements are converted into motivated actions. These behavioral effects are consistent with prior reports. For example, a study found that optogenetic perturbation of cerebellar activity reduces voluntary whisker movements during the photostimulation period and leads to a subsequent rebound in whisking behavior (Proville et al., 2014). We did not assess the circuit effect of autonomous PC photostimulation during behavior. Therefore, we cannot draw any conclusions between the precise pattern of induced activity and motor-event outcomes. However, PC photostimulation produces both direct increases and indirect decreases in simple spiking in the PC ensemble and can drive complex-spike-like bursts of activity (Bonnan et al., 2021; Tsutsumi et al., 2020).

Optogenetic perturbation of PC activity also produced deficits in motor-action timing. Specifically, we found that the perturbation delayed the subsequent peak rate of licking that corresponded to the anticipated time of water-reward availability. Thus, a delay or discoordination in the transition to movement initiation, which is normally signaled by ramping PC activity, disrupts and/or delays the remaining motor plan, leading to a mistimed action. Importantly, the induced behavioral delay only occurred when the time gap between the end of the perturbation and licking onset was less than a second. Such a sub-second scale is consistent with regimes of cerebellar function, such as the timing dependence of synaptic plasticity induction at parallel-fiber-to-PC synapses and motor learning (Chen and Thompson, 1995; Raymond and Lisberger, 1998; Safo and Regehr, 2008; Suvrathan et al., 2016). Optogenetic perturbation of PC activity also delays the onset of sensory-cue-driven licking (Tsutsumi et al., 2020), indicating a consistent effect across movement types. In summary, PC activity both represents and causes motor-event transitions, influencing the coordination of explicitly timed, volitional movements to improve the temporal consistency across repeat cycles of goal-directed action.

## Materials and Methods

### Animals

All animal procedures were performed following protocols approved by the Institutional Animal Care and Use Committee at the Max Planck Florida Institute for Neuroscience. Adult animals (>10 weeks) from the following strains of mice were used in this study: C57/Bl6 (5f, 3m), Pcp2-Cre (1f, 2m), Pcp2-Cre crossed with Ai27 (5f, 8m), and nNOS-ChR2 (1f, 2m) (Kim et al., 2014; Madisen et al., 2012; Zhang et al., 2004). These mice were maintained on a 12 h light-dark cycle with *ad libitum* access to food and were provided a running wheel for enrichment.

### Surgical Procedures

For all surgeries, we used isoflurane for anesthesia (1.5-2%). A heating pad with biofeedback control provided body temperature maintenance. Buprenorphine (0.35 mg/kg subcutaneous), carprofen (5 mg/kg subcutaneous), and a lidocaine/bupivacaine cocktail (topical) were used for pain control. To restrain the head during behavioral experiments, a small stainless-steel post was attached to the skull. Mice had a cranial window installed over the left lateral cerebellum, above a region that included portions of the Crus I and Crus II lobules (Gaffield et al., 2016). For some optogenetic experiments, optical fibers (MFC_400/430-0.48_MF1.25_FLT, Doric Lenses, Quebec, Canada) were instead installed bilaterally over Crus II (3.5 mm lateral, 2.2 mm caudal from lambda). All implants were fixed in position using Metabond (Parkell, Edgewood, NY). For calcium imaging experiments, adeno-associated virus (AAV)1 containing the genetically encoded calcium indicator GCaMP6f (Chen et al., 2013) under control of the Pcp2 promoter (Nitta et al., 2017) was injected into the area under the craniotomy to transduce Crus I and II PCs. All mice were given at least 7 days to recover from surgery before beginning behavioral training and/or experimentation.

### Behavioral Procedures

For the interval timing task, mice were held under head-fixation in a custom-built apparatus consisting of a metal tube (1” diameter) in which the mice rested comfortably and a metal water port that was placed in front of their mouths. This port was calibrated to allocate 4 μl of water per dispensed droplet as determined by the open time of a solenoid valve (INKA2424212H, Lee Company, Westbrook, CT). This apparatus was housed inside a sound-insulated and light-protected enclosure. The water valve was located outside of the enclosure to prevent the mice from hearing it open and close. Licks were detected using a simple transistor-based lick circuit connecting the metal tube to the metal water port. This circuit closed when the tongues of the mice contacted the water port. The apparatus was controlled by a BPod state machine (Sanworks, Rochester, NY) installed on a Teensy 3.6 microcontroller (SparkFun Electronics, Boulder, CO) combined with custom-written codes (Matlab, MathWorks, Natick, MA).

Thirst was used to motivate behavioral performance. To achieve this, mice underwent water restriction with daily water intake limited to 1 ml with frequent monitoring to confirm the lack of any adverse health consequences (Guo et al., 2014). Initial sessions of behavioral training consisted of a block of 50 consecutive water-rewarded trials to reinforce the target time interval. Thereafter, sessions consisted of a trial structure compromising 80% rewarded and 20% unrewarded trials that were randomly distributed. Only a single time interval was used during a session. Mice typically completed 250-300 trials per session, with only a single session per day, and were considered well trained when they consistently initiated licking bouts prior to water delivery, and they accurately anticipated the water delivery time on unrewarded trials (peak lick rate within 0.5 s of the expected time of water allocation). Most mice reached this criterion level after 10-15 training sessions.

### Electrophysiology

The cerebellum was accessed for electrophysiology recording through a previously prepared craniotomy that exposed portions of both the Crus I and II lobules. A glass coverslip was cemented over this area to protect the brain post-surgery while the animals were trained for the behavior. On the day of recording, and under light isoflurane anesthesia, the coverslip was removed by drilling away any Metabond that secured it in place. A silver wire was then place into the craniotomy site to provide a ground signal. At least 45 minutes after recovery from anesthesia, the mouse was transferred to the behavioral apparatus and the exposed cerebellum and the ground wire were covered with a saline solution. A silicon probe (A1×32-Poly3-5mm-25s-177, Neuronexus, Ann Arbor, MI) was then slowly inserted into the brain at a few microns per second under the control of a motorized micromanipulator (Mini, Luigs and Neumann, Ratingen, Germany). The silicon probe was connected to an amplifier (RHD2132, Intan Technologies, Los Angeles, CA) and read out by a controller interface (RHD200, Intan Technologies) with a sampling rate of 20 kHz. Commercial software (Intan Technologies) was used for data acquisition. For electrophysiology experiments combining optogenetics, laser light was delivered directly onto the exposed brain using a patch cable that was carefully positioned to minimize any induced artifacts in the recordings while still targeting the proper region of the cerebellar cortex. A copy of the electrical signal driving the light stimulus was recorded to ensure proper registration to the electrophysiological and behavioral recordings.

### Calcium Imaging

All two-photon imaging experiments were performed as previously described (Gaffield et al., 2016; Gaffield et al., 2019). Briefly, a custom-built, movable-objective microscope acquired continuous images at ~30 frame/s using an 8 kHz resonant scan mirror in combination with a galvanometer mirror (Cambridge Technologies, Bedford, MA). A 16x, 0.8 NA water immersion objective (Olympus, Tokyo, Japan) was used for light focusing and collection. The lens was dipped in an immersion media of diluted ultrasound gel (1:10 with distilled water) that was applied to the cranial window the day of recording. The microscope was controlled using ScanImage software (Vidrio Technologies, Ashburn, VA). GCaMP6f was excited with pulsed infrared (900 nm) light from a Chameleon Vision S laser (Coherent, Santa Clara, CA) with an output power of <30 mW from the objective.

### Optogenetics

In optogenetic experiments, stimulation of ChR2 was driven by a 473 nm continuous-wave laser (MBL-F-473-200mW; CNI Optoelectronics, Changchun, China). An acousto-optical modulator (MTS110-A3-VIS controlled by a MODA110 Fixed Frequency Driver; AA Opto-Electronic, Orsay, France) modulated the laser power to produce brief pulses of light during experimental procedures (40 Hz, 5 ms). For this, the laser light was directed into a patch cable (BFYL2F01; Thorlabs, Newton, NJ) that was either directly placed over the installed cranial window (unilateral stimulation) or connected to the implanted optical fibers (bilateral stimulation). For unilateral stimulation of either Crus I or Crus II, the output of the patch cable was set to 15 mW and the surrounding area was covered with black foam to limit visibility to the mouse. For bilateral stimulation experiments, the light output was ~2.75 mW; the ceramic connectors (ADAL1; Thorlabs) were covered in black heat shrink tubing, and then covered again with small black tubes to limit light visibility to the mouse. All optogenetic experiments included an overhead 470 light emitting diode (M470L4; Thorlabs) that was continuously directed at the mice to mask the light flashes used for optogenetic stimulation.

### Data Analysis

Licking behavior was analyzed by calculating the lick rate as the inverse of the inter-lick interval. To quantify correlations between lick rate and the simple spiking rate in PCs, the mean lick rate for each trial was sorted into 1 s bins. Only trials in which the first lick occurred at least 2 s into the trial were included to ensure enough pre-lick electrophysiological data for analysis. Similarly, only trials in which the last lick had at least 1 s of post-lick electrophysiological data were included in the analysis. For optogenetic perturbation experiments, the lick variance was normalized to the pre-stimulus control condition for each session. To calculate the relative change in lick rate after the optogenetic-stimulus-induced rebound in movement, we computed the difference between the lick rates in control trials and photostimulation trials, and then divided by the control rate.

Silicon probe data were sorted automatically by the Kilosort algorithm (Pachitariu et al., 2016), followed by manual curation using Phy2 software (https://github.com/cortex-lab/phy) into 202 unique clusters. In a first-pass analysis, PC units (n = 48) were unambiguously identified by accompanying complex spikes in the recordings (*Figure 2-figure supplement 1A*). We also confirmed that these units had the simple spiking properties expected for PCs (i.e., firing rate and regularity) (Van Dijck et al., 2013). There were an additional 56 units with similar simple spiking responses, but complex spikes could not be clearly discerned in their activity recordings. Noticeably, spiking in unambiguously identified PC units was clearly represented in many nearby channels of the silicon probes (*Figure 2-figure supplement 1C*). Therefore, we calculated the mean peak spike size in each channel and counted the number of channels with a peak significantly above the noise (~30 μV). This approach determined that the channel count for unambiguously identified PCs tended to be quite high (i.e., ≥ 7), likely due to the large size of PCs relative to the spacing of the electrode pads on the silicon probes (*Figure 2-figure supplement 1C,D*).

To test whether channel count representation could be used to classify PCs, we compared the spike distribution of unambiguously identified PCs in each channel with those of another abundant cerebellar cell type, molecular layer interneurons (MLIs). We did not choose to examine granule cells, another abundant cell type, in this comparison because granule cells typically have very different behavior-evoked firing properties than PCs (Powell et al., 2015). In addition, due to the low impedance of the electrode pads used on silicon probes, the activity of granule cells is generally not detectable in extracellular electrophysiological recordings. To positively identify MLI units in our recordings, we used an opto-tag strategy whereby ChR2-expressing MLIs in nNOS-ChR2 mice (Kim et al., 2014) had short-latency responses to light pulses (*Figure 2-figure supplement 1B*; recordings were not obtained from these mice during behavior). In comparison to unambiguously identified PC units, spiking in identified MLI units was detected in only a few channels (*Figure 2-figure supplement 1C-E*). Therefore, a high representation of spiking activity in many channels (≥ 7) could successfully classify PCs. Using this criterion to identify additional putative PCs, we included another 41 units, from the 56 showing similar spiking response characteristics, for a total of 89 in our analysis.

We manually examined the data to classify PCs into groups based on how their simple spiking activity was tuned to different types of motor-event transitions during the behavioral task. PCs were classified as activated during the first lick in a bout if their simple spiking rate showed three consecutive increases in 100 ms time bins that began 1 s before that first lick and ended 0.5 s after that same first lick. The ramp onset time was the time bin of the first of those rate increases. PCs were classified as activated during the last lick in a bout based on the mean simple spike rate during the 300 ms preceding the last lick, with a z-score > 1 as a classification criterion for being last-lick related. Negatively modulated PCs were classified using the same criterion except we used negative values instead. In some cases, individual PC recordings were noisy enough that our rigid classification may have missed some responding cells, so we expect our results to represent a lower-bound estimate of tuning specificity.

Calcium imaging data were analyzed using a standard procedure (Gaffield et al., 2016). In brief, individual PC dendrites were identified using independent component analysis (Hyvarinen, 1999). Calcium events were then extracted from the raw fluorescence traces using an inference algorithm (Vogelstein et al., 2010). Regions of interest (ROIs) in the imaged areas of Crus I and II were considered water-responsive if a peak in activity was observed in the total PC average for that region at the time of water delivery. The ramp onset time was determined when the peak event rate reached > 3 standard deviations above the mean, in 100 ms time bins, beginning 1 s before and up to 0.5 s after the first lick in a lick bout. To quantify calcium event activity prior to the last lick, we used a similar criterion, but examined event rate activity at least 2 s post water delivery in rewarded trials as well as at the end of licking in water-omission trials.

In the figures, shaded areas in activity plots indicate SEM range; error bars are also represented as SEM. Statistical values were calculated using GraphPad (Prism, San Diego, CA) with significance indicated by p-values below 0.05.

## Acknowledgements

We thank Samantha Amat for laboratory assistance and the GENIE program (Janelia Research Campus, including Drs. Jayaraman, Kerr, Kim, Looger, and Svoboda) for freely providing GCaMP6f to the neuroscience community. This work was supported by National Institutes of Health Grants NS083894 and NS105958 (J.M.C.) and the Max Planck Florida Institute for Neuroscience.

## Supplemental Figures

**Figure 2-figure supplement 1.**
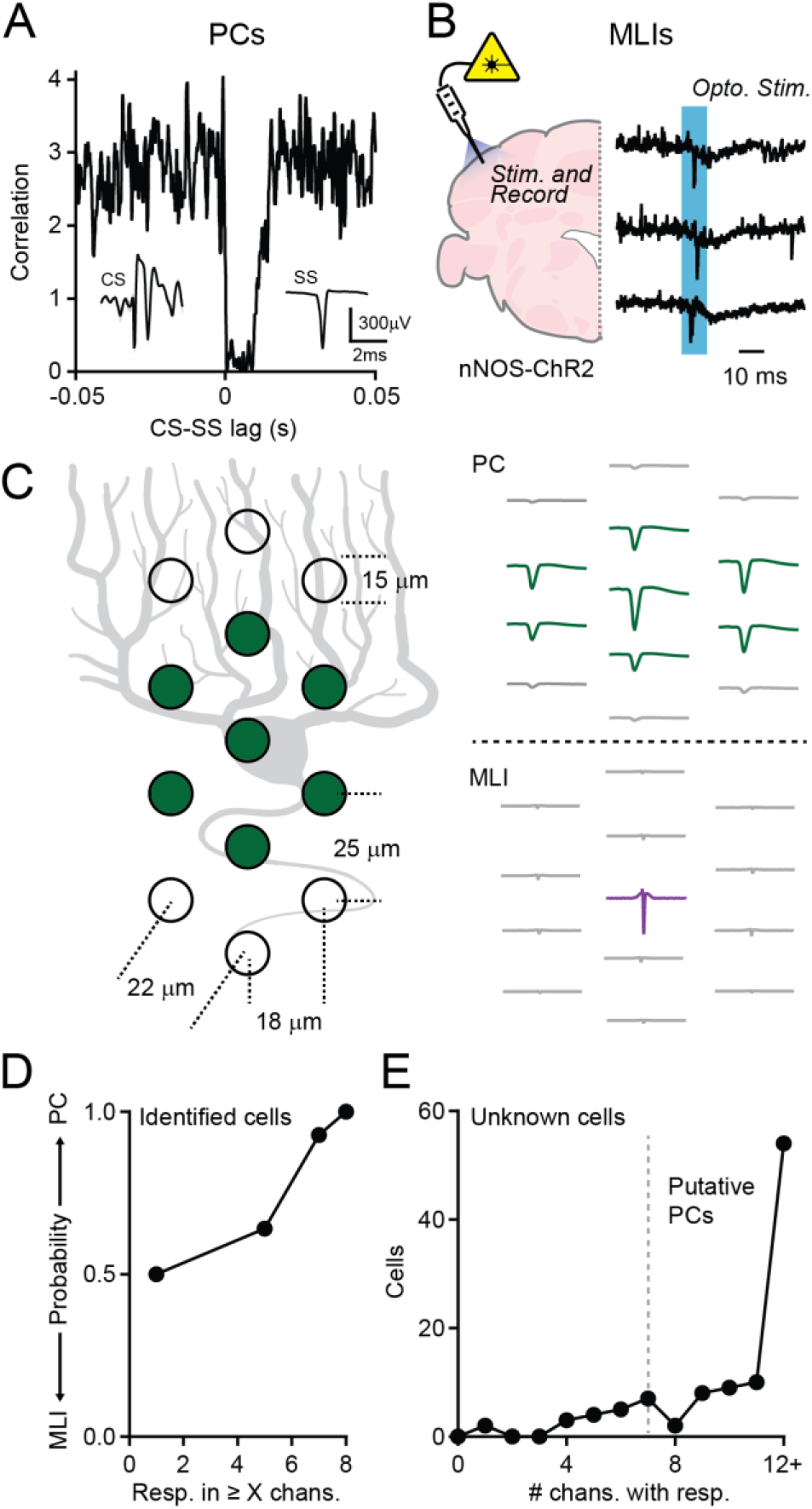
Identification of PCs in silicon probe data. A. PCs were unambiguously identified by the appearance of both complex spikes (left inset) and simple spikes (right inset) in the unit recordings. The correlation plot of these two spike types revealed the characteristic complex-spike-induced pause in simple spike firing. Data are from an individual PC. B. MLIs were identified using an opto-tag strategy. Specifically, ChR2-expressing MLIs showed short-latency spiking responses to brief optogenetic stimuli. Example traces show evoked activity in three different MLIs from the same mouse with the photostimulus period indicated in blue. C. Left: Illustration of the arrangement of electrode pads on the silicon probe overlaid on a prototypic PC. Top Right: Mean spikes recorded in each electrode channel for an example PC. Bottom Right: Mean spikes recorded in each electrode channel for an example MLI. The purple trace is the only channel with a response. D. Plot of the probability that cells whose spikes were observed in at least X channels had a confirmed identity as either a PC (n = 23 cells) or an MLI (n = 25 cells, total of 3 mice). E. Histogram of all potential PCs, based on simple spike firing characteristics (n = 104), and the number of channels with detectable spikes for each cell. Based on the analysis in panel D, all cells with spikes in 7 or more channels were classified as PCs.

**Figure 7-figure supplement 1.**
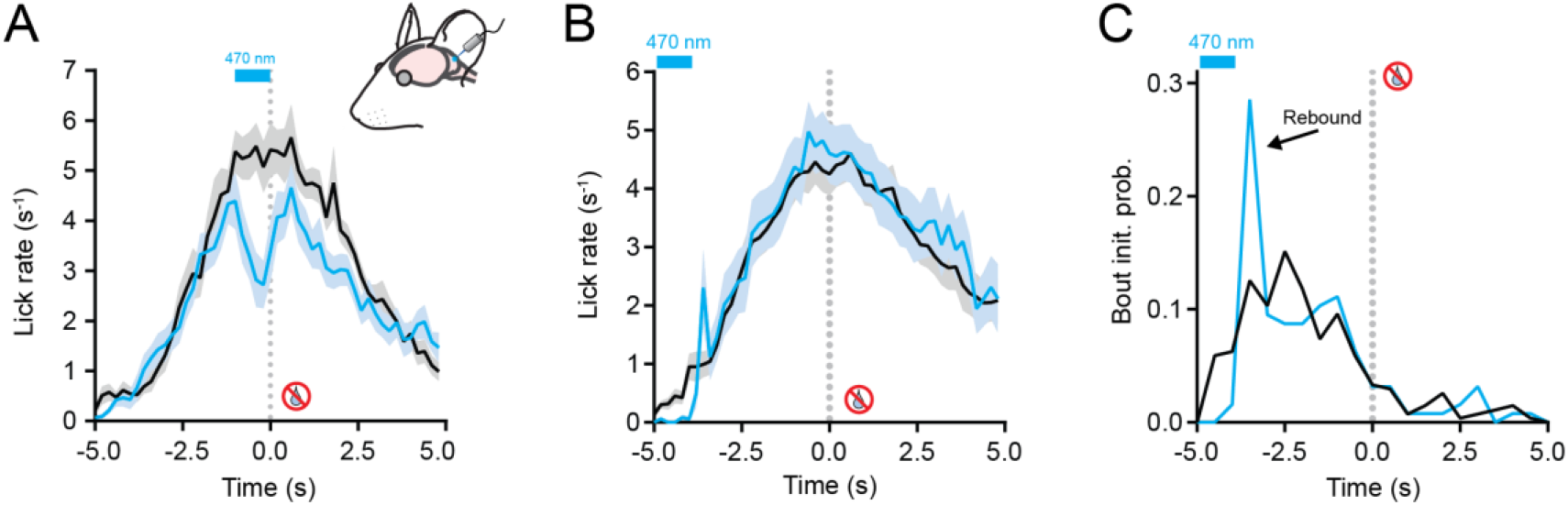
Unilateral optogenetic stimulation of PCs during the task. A. Unilateral optogenetic stimulation was delivered to ChR2-expressing PCs in left Crus I or Crus II during the period of peak licking in water-omission trials. Comparison of average licking rates for stimulation trials (blue) as well as interleaved control trials (black) (n = 12 sessions, 4 mice). B. As in panel A but with the photostimulus timed to the period of earliest lick-bout initiations (n = 20 sessions, 7 mice). C. Histogram of the lick bout initiation times. A clear rebound in lick probability is apparent immediately after the unilateral photostimulation ended.

**Figure 8-figure supplement 1.**
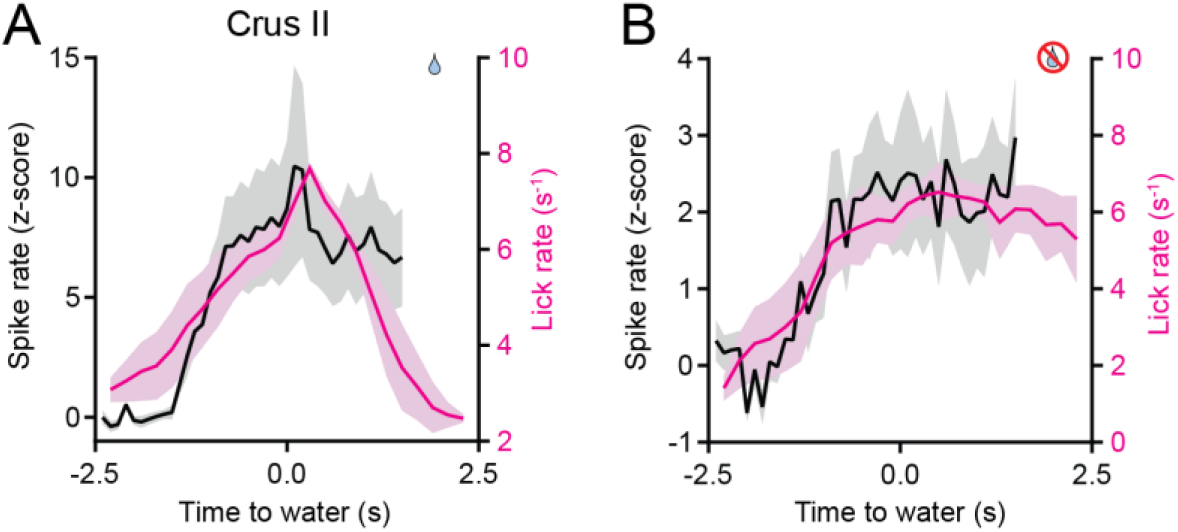
Modulation of PC simple spiking during a short-interval timing task. A. Simple spike rates from PCs (n = 17 from 3 mice; black) and the corresponding lick rates (pink lines) are shown for Crus II recordings during water-rewarded trials with a 5 s timing interval. B. Same as panel A for unrewarded, water-omission trials.

